# Snake deltavirus utilizes envelope proteins of different viruses to generate infectious particles

**DOI:** 10.1101/698514

**Authors:** Leonora Szirovicza, Udo Hetzel, Anja Kipar, Luis Martinez-Sobrido, Olli Vapalahti, Jussi Hepojoki

## Abstract

Hepatitis D virus (HDV) is the only known human satellite virus, and it requires hepatitis B virus (HBV) to form infectious particles. Until 2018 HDV was the sole representative of the genus *Deltavirus*. The subsequent identification of HDV-like agents in non-human hosts indicated a much longer evolutionary history of deltaviruses than previously hypothesized. Interestingly, none of the HDV-like agents were found in co-infection with an HBV-like agent suggesting that these viruses use different helper virus(es). Here we show, using snake deltavirus (SDeV), that HBV represents only a single example of helper viruses for deltaviruses. We cloned the SDeV genome into a mammalian expression plasmid, and by transfection could initiate SDeV replication in cultured snake and mammalian cells. By superinfecting SDeV infected cells with arenaviruses, or by transfecting different viral surface proteins, we could induce infectious SDeV particle production. Our findings indicate that deltaviruses can likely use a multitude of helper viruses or even viral glycoproteins to form infectious particles. Persistent and recurrent infections would be beneficial for the spread of deltaviruses. It seems plausible that new human and veterinary disease associations with deltavirus infections will be identified in the future.

## INTRODUCTION

Viroids found in higher plants are the smallest known infectious agents comprised of only circular RNA^1^. After the discovery of viroids in 1971^2^, hepatitis D virus (HDV) was described in 1977^3^ as the first human pathogen with an RNA genome resembling viroids. Alike viroids, the circular genome of HDV forms secondary structures by self-complementarity, although the genome of HDV is roughly four times bigger^4^. Viroids also replicate by the rolling circle mechanism and possess ribozyme activity^4^. However, unlike viroids, HDV encodes a functional protein and it also requires hepatitis B virus (HBV) as a helper virus^4^. Until recently, HDV has only been found in humans. Then, in 2018, HDV-like sequences were reported from two non-human hosts^5, 6^. The findings challenged the view on the origin and evolution of HDVs within their human host^7^.

HDV is unique among animal viruses, and forms the genus *Deltavirus*, which has not been assigned to a family^8^. The negative-sense single-stranded RNA genome of HDV is approximately 1.7 kb, circular and highly self-complementary, due to which it forms unbranched rod-like structures^9, 10^. During replication both genomic and antigenomic viral RNA are found in the infected cells^11^. The only conserved open reading frame (ORF) of HDV is in antigenomic orientation and encodes the hepatitis delta antigen (HDAg, used for HDV)^9^. Both RNA strands possess a ribozyme activity responsible for self-cleavage^12^. The ribozyme was initially speculated to also mediate the ligation to form the circular genome^13^, but later studies indicated involvement of host enzymes^14^. As HDV only encodes HDAg, it cannot form infectious particles without a helper virus^15^. The discovery of HDV in liver specimens of HBV positive individuals directly associated HDV with HBV, as a satellite virus^3^. Later studies demonstrated the transmissible and pathogenic nature of HDAg^16^, and that HDV relies on the envelope proteins of HBV to form infectious particles^15^. Although helper virus is required for producing infectious particles, the rolling-circle replication^11^ proceeds independently of the helper virus^17^, mediated by host RNA polymerase II^18^. During the viral life cycle two different forms of HDAg, small-(S-) and large (L-) HDAg, are produced^19^. The HDAg ORF encodes the S-HDAg, and a base transition in the amber stop codon results in the elongation of the protein at the carboxy terminus by 19 additional amino acids, thus giving rise to the L-HDAg^20^. The two antigen forms play highly diverse roles in the viral life cycle, e.g. S-HDAg promotes while L-HDAg inhibits viral replication^19, 21^. In fact, L-HDAg also suppresses the expression of HBV proteins, and acts in the assembly of infectious particles^22^.

Due to the symbiotic relationship with HBV, HDV infection is acquired either via co-infection with HBV or via superinfection of a chronically HBV infected individual^23^. The disease outcome varies greatly and depends on the mode of HDV infection^24^. Co-infection often results in acute hepatitis, which tends to be self-limited, whereas superinfection can lead to a fulminant hepatitis which in many cases becomes chronic, ultimately leading to liver cirrhosis^25^. In these patients, the risk of liver failure or development of subsequent hepatocellular carcinoma is high^25^. However, the disease is also influenced by the HDV genotype^24^; of these, eight are currently known^26^. While HDV-1 occurs worldwide, the other genotypes have a specific geographic distribution^26^. Interestingly, a very recent report described HDV to be capable of producing infectious particles utilizing envelope glycoproteins of several viruses^27^.

The report on discovery of a HDV-like agent in birds^5^ urged us to publish our observation of a similar, yet genetically distant agent in snakes^6^. Both the avian and snake deltaviruses (AvDV and SDeV, respectively) possess a negative-sense, highly self-complementary circular RNA genome including ribozymes^5, 6^. The findings complemented one another in that no hepadnaviral sequences were identified in the samples^5, 6^. Moreover, we could demonstrate the presence of both viral RNA and snake delta antigen (SDAg) in several tissues, indicating that the replication of SDeV is not restricted to the liver^6^, which likely holds true also for AvDV. To further support the idea that the evolutionary path of HDV is much longer than initially envisioned, more HDV-like agents were identified in fish, amphibians and invertebrates in early 2019^28^. Similar to AvDV and SDeV, the newly found deltaviruses were not associated with hepadnavirus infection^28^. The new findings have raised the question if deltaviruses are indeed dependent on hepadnavirus co-infection. The authors of the report on AvDV identified sequences matching to influenza A virus genome in the same samples^5^, and similarly we could demonstrate replication of both reptarenaviruses and hartmaniviruses in the snakes with SDeV^6^. These findings led us to hypothesize that deltaviruses have evolved to use enveloped viruses that cause persistent infection as their helpers. The aim of our study was to experimentally demonstrate that SDeV can indeed form infectious particles utilizing the envelope proteins of other than hepadnaviruses.

## RESULTS

### Isolation of SDeV from the brain of an infected snake

We originally identified SDeV when performing a metatranscriptomic analysis of a brain sample from a snake with central nervous system signs^6^. Subsequent RT-PCR screening demonstrated the presence of SDeV in multiple tissues including liver and blood. We then used metatranscriptomic analysis to look for traces of HBV-like virus in blood and liver, but instead retrieved genomes of co-infecting reptarena- and hartmaniviruses. Since hepadnaviruses are hepatotrophic, we reasoned that successful isolation from the brain sample would indicate that SDeV could utilize arenaviruses for infectious particle formation. We inoculated cultured boid kidney cells (I/1Ki) with the infected brain homogenate, and at 15 days post infection (dpi) analyzed the cells by immunofluorescence (IF) staining. We used affinity purification to produce anti-SDAg and anti-NP (reptarenavirus nucleoprotein, the main antigen present in infected cells) reagents, which we directly labelled by AlexaFluor488 or AlexaFluor594 dyes (anti-SDAg-AF488, anti-SDAg-AF594, anti-NP-AF488, and anti-NP-AF594). As demonstrated in Figure 1, the reagents produce hardly any background, and can be used for co-staining of SDeV and reptarenaviruses. Figure 1 further demonstrates that we were able to induce SDeV infection via inoculation with the infected brain homogenate. We further tested whether the infected cells produce progeny virions by titrating the supernatant collected from the infected cells at 7 dpi on clean I/1Ki, and recorded titers of 4.0*10^3^ fluorescent focus-forming units/ml (fffus/ml).

**Figure 1.**
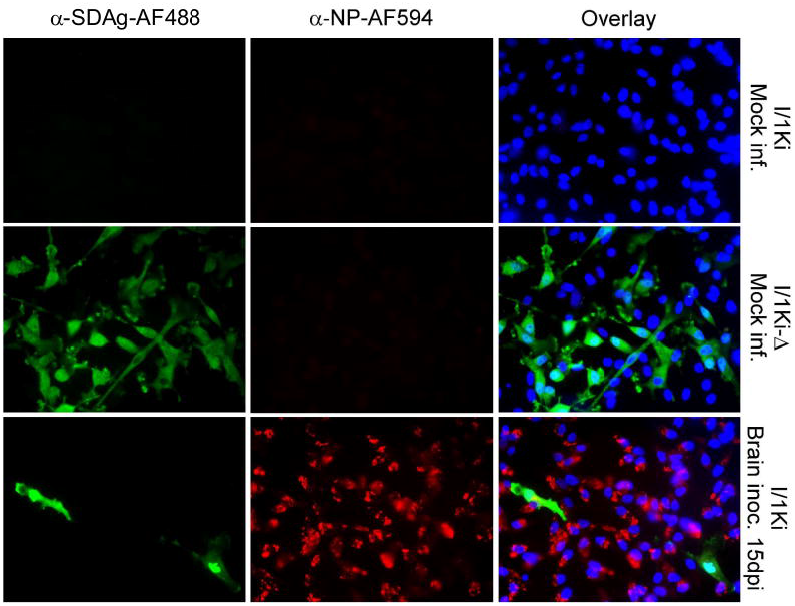
Isolation of SDeV from the brain of an infected snake using cultured boid kidney cells (I/1Ki). Mock-infected I/1Ki (top panels), mock-infected I/1Ki-Δ (middle panels), and brain homogenate inoculated I/1Ki cells (bottom panels) were stained for SDAg (α-SDAg-AF488, left panels, green), reptarenavirus nucleoprotein (α-NP-AF594, middle panels, red), and Hoechst 33342 was used to visualize the nuclei, the panels on right show an overlay of the three images. The images were taken at 400x magnification using Zeiss Axioplan 2 microscope.

### Transfection of cultured cells with SDeV constructs initiates replication of the virus

After successfully isolating SDeV in cell culture, we wanted to study SDeV replication without co-infecting viruses. As replication of HDV occurs via rolling circle replication^11^, we decided to generate expression constructs with multiple copies of the SDeV genome, an approach successfully applied for HDV reverse genetics^17^. We ordered a synthetic gene comprising two copies of the SDeV genome, and subcloned the insert in genomic (pCAGGS-SDeV-FWD) and antigenomic (pCAGGS-SDeV-REV) orientation into the mammalian expression vector pCAGGS/MCS. The expression constructs are schematically presented in Figure 2. We then used the generated constructs to transfect *B. constrictor* (I/1Ki) and African green monkey (Vero E6) kidney cells. IF staining of the transfected cells at 1, 2, 3, and 4 days post transfection demonstrates that SDAg can be detected from day 1 onwards (Figure 3A). The result further indicates that SDAg is expressed in cells transfected with either pCAGGS-SDeV-FWD or pCAGGS-SDeV-REV. Since the SDAg is encoded in the antigenomic orientation, we interpret the detection of SDAg in cells transfected with pCAGGS-SDeV-FWD plasmid as an indication of replication. Interestingly, at 1 to 2 days post transfection the SDAg was predominantly found in the cytoplasm when expressed from the pCAGGS-SDeV-REV plasmid, whereas transfection with the pCAGGS-SDeV-FWD plasmid resulted in most of the SDAg presenting in nuclear localization. At four days post transfection, however, SDAg was mostly detected in the cytoplasm for both constructs. We also analyzed the transfected cells by western blotting (WB) and could demonstrate an increasing amount of SDAg during the four days of transfection (Figure 3B).

**Figure 2.**
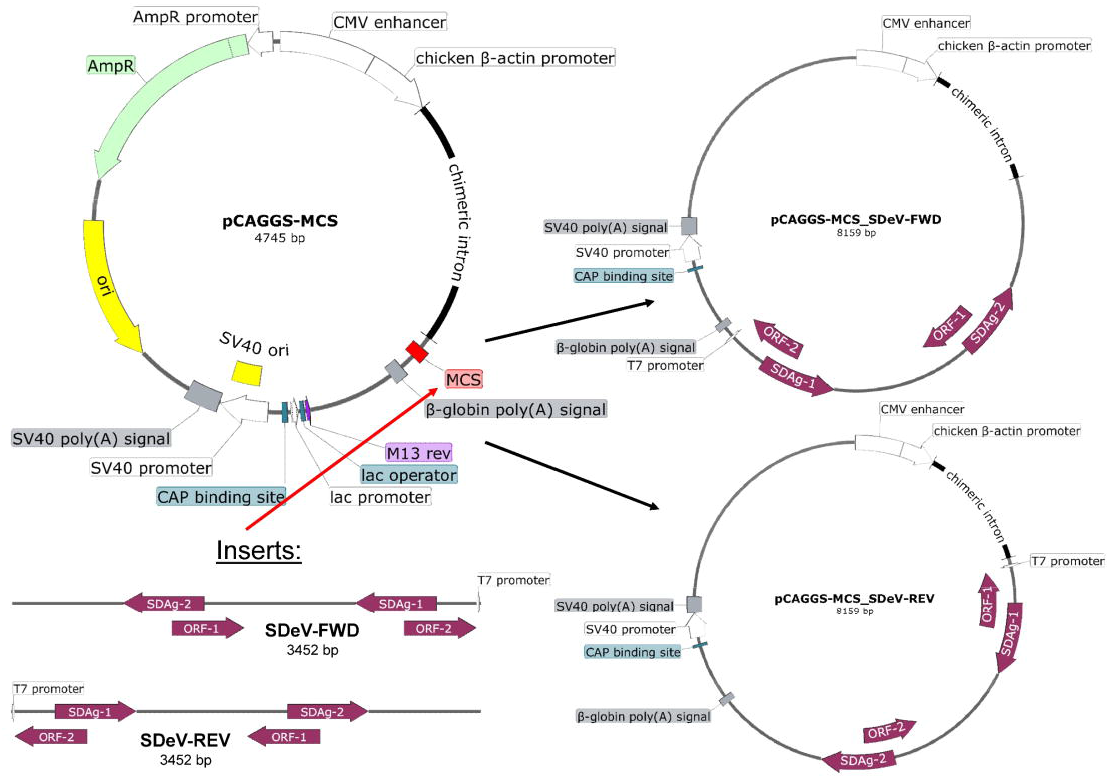
A schematic representation of the construction of the SDeV recombinant expression plasmids. The vector backbone (pCAGGS-MCS) is shown top left and the final constructs are shown on the right. The inserts (bottom left) containing two copies of the SDeV genome were cloned into the MCS (multiple cloning site) as indicated by the red arrow. The inserts contain two copies of the SDAg (SDAg-1 and −2), open reading frame (ORF-1 and −2) for an unknown protein product and T7 promoter. Blunt end cloning was used to obtain constructs (shown on the right) with the insert in genomic (pCAGGS-SDeV-FWD) or antigenomic (pCAGGS-SDeV-REV) orientation.

**Figure 3.**
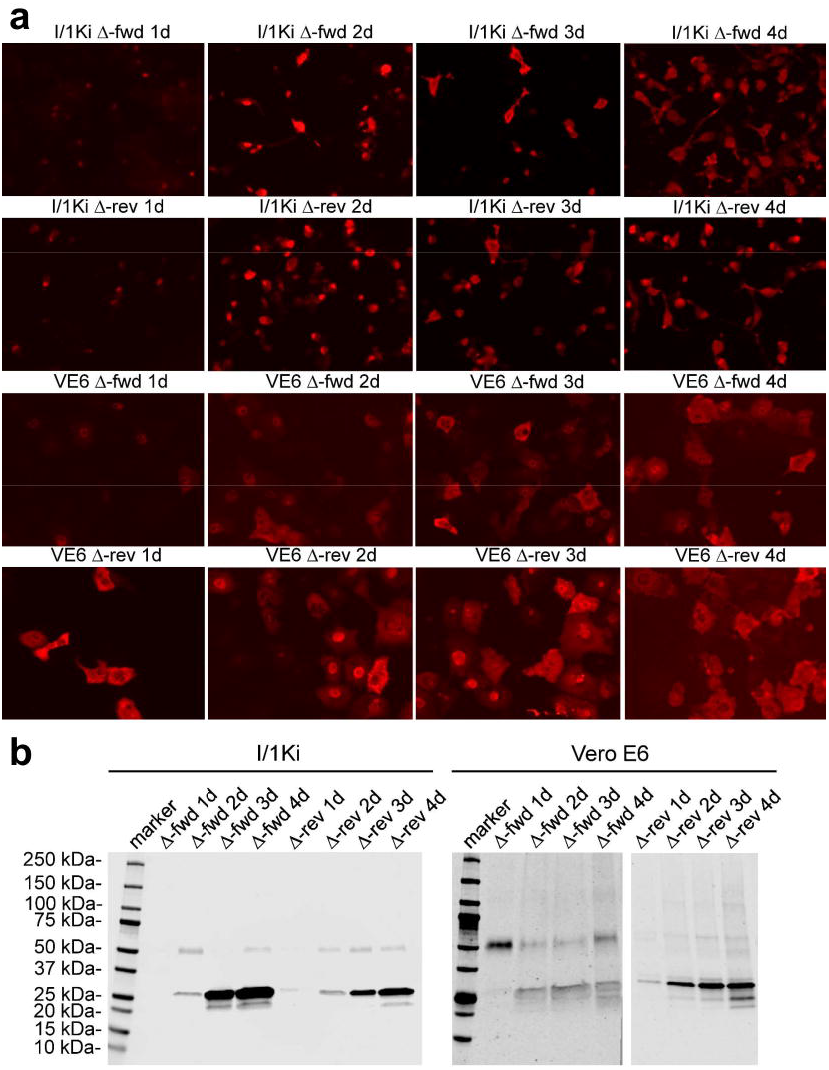
Transfection of I/1Ki and Vero E6 cells with pCAGGS-SDeV-FWD and pCAGGS-SDeV-REV constructs results in SDeV replication. **a)** I/1Ki (top) and Vero E6 (bottom) cells transfected with Δ-fwd (pCAGGS-SDeV-FWD) and Δ-rev (pCAGGS-SDeV-REV) were stained for SDAg (anti-SDAg antiserum 1:7,500 and Alexa Fluor 594-labeled donkey anti-rabbit immunoglobulin 1:1,000) at 1, 2, 3, and 4 days post transfection (from left to right). The images were taken at 400x magnification using Zeiss Axioplan 2 microscope. **b)** Western blot of I/1Ki (left panel) and Vero E6 (right panel) cell pellets at 1, 2, 3, and 4 days post transfection with Δ-fwd and Δ-rev constructs. Precision Plus Protein Dual Color Standards (Bio-Rad) served as the marker, and the results were recorded using Odyssey Infrared Imaging System (LI-COR).

### Replication of SDeV in human and snake cells

After demonstrating that replication of SDeV can be initiated in both boid and monkey kidney cells by transfecting the pCAGGS-SDeV-FWD plasmid, we wanted to test if SDeV replication would also occur in human cell lines. We transfected human lung carcinoma (A549), hepatocyte carcinoma (Hep G2), cervical cancer (HeLa), and embryonic kidney (HEK293FT) cell lines with the pCAGGS-SDeV-FWD plasmid, and used IF to demonstrate the presence of SDAg. IF staining at five days post transfection showed SDAg in a cytoplasmic localization in all cell lines studied (Figure 4A). In addition, SDAg was also strongly expressed in the nucleus of A549 and Hep G2 cells. To study whether similar differences in the SDAg localization would occur also in snake cells, we transfected boid kidney (I/1Ki and V/1Ki), heart (V/2Hz), liver (V/1Liv), and lung (V/5Lu) cell lines with the pCAGGS-SDeV-FWD plasmid. We performed IF staining five days post transfection where SDAg demonstrated a variable expression pattern depending on the cell line (Figure 4B). However, in all cell lines, SDAg was found both in the cytoplasm and the nucleus. Curiously, the localization appeared to be more nuclear in liver and heart cell lines.

**Figure 4.**
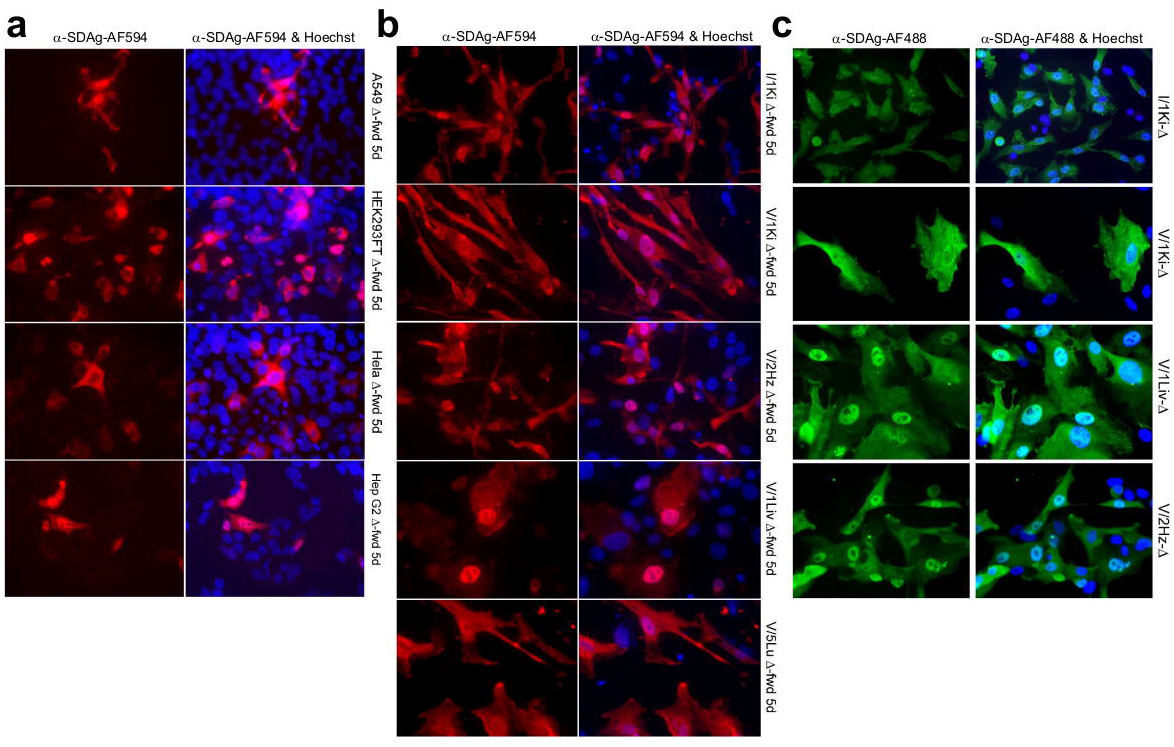
SDeV replicates in human and reptilian cell lines. **a)** A549 (human lung carcinoma), HEK293FT (human embryonic kidney), HeLa (human cervical cancer), and Hep G2 (human hepatocellular carcinoma) (from top to bottom) cells transfected with Δ-fwd (pCAGGS-SDeV-FWD) were stained at 5 days post transfection for SDAg (α-SDAg-AF594, left panels, red) and Hoechst 33342 was used to visualize the nuclei, the panels on right show an overlay. **b)** Boid cell lines I/1Ki (kidney), V/1Ki (kidney), V/2Hz (heart), V/1Liv (liver), and V/5Lu (lung) (from top to bottom) transfected with Δ-fwd (pCAGGS-SDeV-FWD) were stained at 5 days post transfection for SDAg (α-SDAg-AF594, left panels, red) and Hoechst 33342 was used to visualize the nuclei. The panels on right show an overlay. **c)** The transfected boid cells from **b** were allowed to grow, passaged three times, and stained for SDAg (α-SDAg-AF488, left panels, green) and Hoechst 33342 was used to visualize the nuclei. The panels on the right show an overlay. All images were taken at 400x magnification using Zeiss Axioplan 2 microscope.

### Transfection of pCAGGS-SDeV-FWD into boid cells results in persistent SDeV infection

After demonstrating that transfection of cultured cells with pCAGGS-SDeV-FWD induces SDeV replication, we wanted to study the effect of prolonged maintenance of the transfected cells under normal culturing conditions. The extent of the cytopathic effect induced by the initial transfection varied a lot depending on the cell line used, but we allowed the surviving cells to reach confluency. We passaged the cells until they reached a surface area of approximately 150-175 cm^2^, and all except the lung cell line (V/5Lu) revived efficiently after the transfection. We then analyzed the cell lines for the presence of SDAg by IF (Figure 4C). SDAg was found in all cell lines, but the size of the persistently SDeV infected cell population varied between the cell lines. For I/1Ki, V/1Liv, and V/2Hz most cells displayed SDAg indicating active replication, whereas for V/1Ki only 5-10% of the cells were SDAg positive. The localization of SDAg varied between the different cell lines, but most often SDAg was found in both cytoplasm and nucleus, which is similar to what we observed *in vivo*^6^. We labelled the persistently SDeV infected cell lines as: I/1Ki-Δ, V/1Ki-Δ, V/1Liv-Δ, V/2Hz-Δ, and V/5Lu-Δ.

Encouraged by these findings, we tried the same approach for mammalian cells (Vero E6), but based on IF screening this cell line was not able to maintain SDeV infection when cultured at 37 °C. To see if temperature is an influencing factor, we then kept the transfected Vero E6 cells at 30 °C, but temperature had little or no effect on virus replication as judged by the number of cells displaying SDAg.

### Superinfection of I/1Ki-**Δ** cells with reptarenaviruses and hartmaniviruses produces infectious SDeV particles

As we had succeeded in isolating SDeV from the brain of a *B. constrictor* that showed no traces of a co-infecting hepadnavirus, but instead carried several reptarenavirus and hartmanivirus L and S segments^6, 29^, we thus wanted to investigate if the permanently infected cell lines could be superinfected with reptarenaviruses and/or hartmaniviruses. So we incubated I/1Ki-Δ cells with reptarenavirus (University of Helsinki virus-2, UHV-2; and University of Giessen virus-1, UGV-1) or hartmanivirus (Haartman Institute Snake virus-1, HISV-1). Figure 5A shows that I/1Ki-Δ cells can be superinfected with reptarenaviruses (UHV-2 and UGV-1). The localization of HDAg is known to change from nuclear to cytoplasmic during the viral life cycle^30^. While most I/1Ki-Δ cells displayed SDAg in the cytoplasm, some cells showed a granular nuclear SDAg staining, similarly to HDAg in human hepatocytes^31^. However, granules appeared less abundant in the reptarenavirus superinfected I/1Ki-Δ cells (Figure 5A). We then tested if the other permanently deltavirus infected cell lines could be superinfected with reptarenavirus (UGV-1) or hartmanivirus (HISV-1). IF staining shows that superinfection was not efficient in V/1Ki-Δ cells (Figure 5B), however, both V/Liv-Δ (Figure 5C) and V/2Hz-Δ (Figure 5D) could be superinfected with both viruses. The shift of SDAg from the nucleus to the cytoplasm was less clear in the other cell lines tested, and further studies are needed to determine if co-infection actually affects the localization of SDAg.

**Figure 5.**
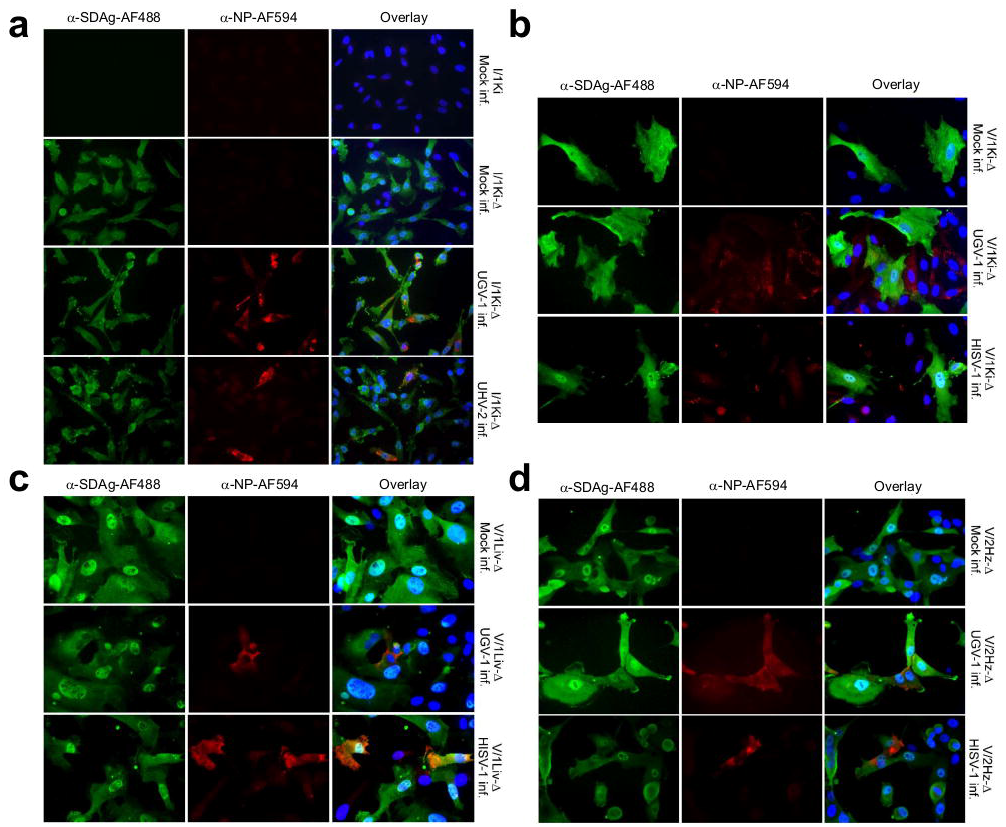
SDeV infected cells can be superinfected with reptarenaviruses (UHV-2 and UGV-1) and hartmanivirus (HISV-1). **a)** Mock-infected I/1Ki cells (boa kidney) and mock-, UGV-1, and UHV-2 infected I/1Ki-Δ cells were stained for SDAg (α-SDAg-AF488, left panels, green), reptarenavirus NP (a-NP-AF594, middle panels, red), and Hoechst 33342 was used to visualize the nuclei, the panels on right show an overlay. **b)** Mock-, UGV-1, and HISV-1 infected V/1Ki-Δ cells (boa kidney) were stained for SDAg (α-SDAg-AF488, left panels, green), reptarenavirus NP (α-NP-AF594, middle panels, except bottom, red) or hartmanivirus NP (anti-HISV NP 1:3,000 and Alexa Fluor 594-labeled donkey anti-rabbit immunoglobulin 1:1,000, middle panel bottom, red), and Hoechst 33342 was used to visualize the nuclei, the panels on right show an overlay. **c)** Mock-, UGV-1, and HISV-1 infected V/1Liv-Δ cells (boa liver) were stained for SDAg (α-SDAg-AF488, left panels, green), reptarenavirus NP (a-NP-AF594, middle panels, red) or hartmanivirus NP (anti-HISV NP 1:3,000 and Alexa Fluor 594-labeled donkey anti-rabbit immunoglobulin 1:1,000), and Hoechst 33342 was used to visualize the nuclei, the panels on right show an overlay. **d)** Mock-, UGV-1, and HISV-1 infected V/2Hz-Δ cells (boa heart) were stained for SDAg (α-SDAg-AF488, left panels, green), reptarenavirus NP (a-NP-AF594, middle panels, red) or hartmanivirus NP (anti-HISV NP 1:3,000 and Alexa Fluor 594-labeled donkey anti-rabbit immunoglobulin 1:1,000), and Hoechst 33342 was used to visualize the nuclei, the panels on right show an overlay. All images were taken at 400x magnification using Zeiss Axioplan 2 microscope.

We then wanted to study whether reptarena- or hartmanivirus superinfection of I/1Ki-Δ cells would induce formation of infectious SDeV particles. We chose to use I/Ki-Δ cells since we have demonstrated that the cell line is permissive for several viruses^29, 32–34^. We inoculated I/1Ki-Δ cells with UHV-2, UGV-1, or HISV-1, collected supernatant up to eight dpi, and analyzed it for infectious particles. We then inoculated a fresh monolayer of clean I/1Ki cells with the supernatants, and used supernatant collected from non-superinfected (mock) I/1Ki-Δ cells as the control. At 2-5 dpi we IF-stained the cells for SDAg and counted the number of fluorescent foci at each time point. The non-superinfected I/1Ki-Δ cells did not produce infectious particles, while cells superinfected with either reptarenaviruses or hartmaniviruses produced infectious SDeV particles (Figure 6A). The production of infectious SDeV particles appeared to be most efficient in HISV-1 infected cells, while UHV-2 infected cells produced the lowest amount of infectious SDeV particles (Figure 6A). The observed difference between the amounts of infectious SDeV particles produced in UHV-2 vs. UGV-1 infected cells might be related to the comparatively lower replication rate of UHV-2 reported in our previous study^29^.

**Figure 6.**
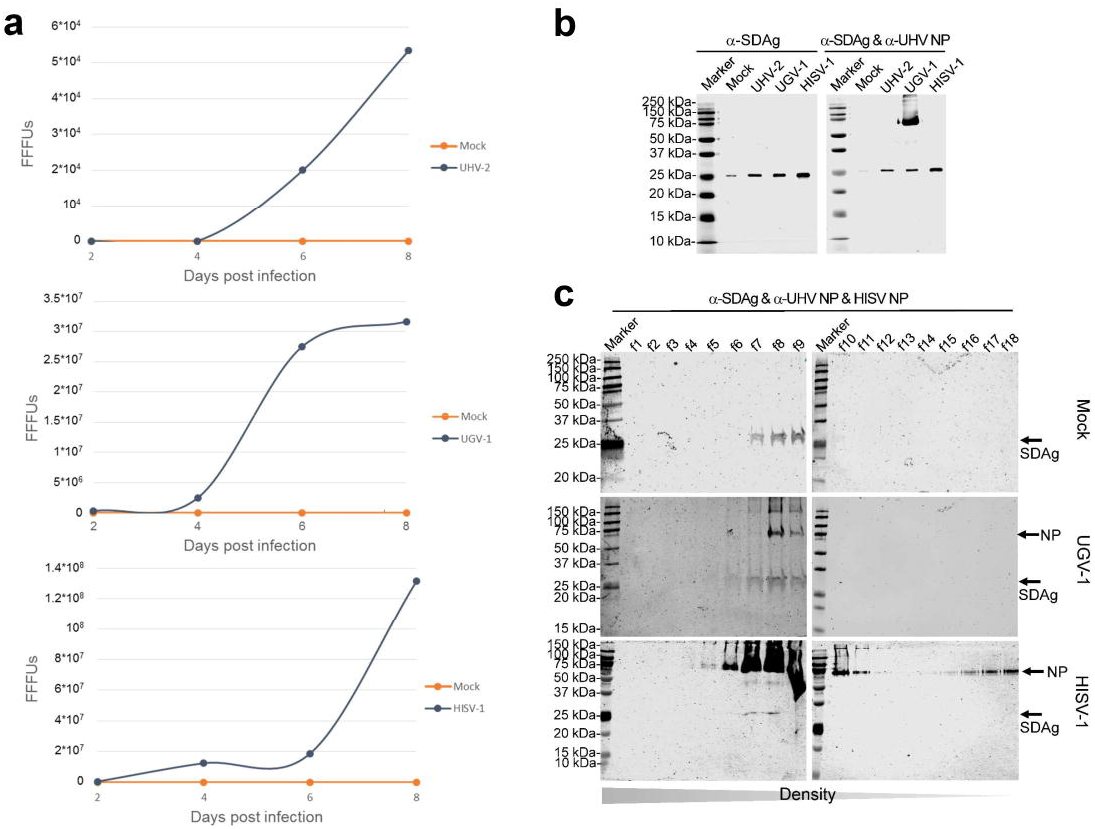
Superinfection of permanently SDeV infected boid kidney cells (I/1Ki-Δ) induces production of infectious SDeV particles. **a)** Supernatant collected from mock-, UHV-2 (top), UGV-1 (middle) and HISV-1 (bottom) superinfected I/1Ki-Δ cells at 2, 4, 6, and 8 days post infection (dpi) were titrated on clean I/1Ki cells. The y-axis shows the number of fluorescent focus forming units (fffus) per ml of culture medium. **b)** Supernatants collected from mock-, UHV-2, UGV-1, and HISV-1 superinfected I/1Ki-Δ cells were pelleted by ultracentrifugation and analyzed by western blotting. The left panel shows anti-SDAg staining and the panel on the right shows anti-SDAg and anti-reptarenavirus NP staining. **c)** Pelleted supernatants collected from mock- (top panels), UGV-1 (middle panels), and HISV-1 (bottom panels) superinfected I/1Ki-Δ cells were subjected to density gradient ultracentrifugation, and the fractions collected from the bottom of the tubes were analyzed by western blotting using anti-SDAg and anti-reptarenavirus NP staining (for mock and UGV-1) or anti-SDAg and anti-hartmanivirus NP (for HISV). The arrows indicate the location of SDAg and reptarenavirus/hartmanivirus NP. Precision Plus Protein Dual Color Standards (Bio-Rad) served as the marker for both **b** and **c**, and the results were recorded using Odyssey Infrared Imaging System (LI-COR).

We then wanted to try purification of the SDeV particles and used ultracentrifugation to pellet the particles secreted from mock, UHV-2, UGV-1, and HISV-1 infected I/1Ki-Δ cells; WB served to detect SDAg in the obtained pellets. The results show that more SDAg could be pelleted from the supernatants of reptarenavirus and hartmanivirus superinfected I/1Ki-Δ cells than from the mock-infected I/1Ki-Δ cells (Figure 6B). Interestingly, also the mock infected cells released particles containing SDAg, however, these particles are non-infectious as described above (Figure 6A). We used density gradient ultracentrifugation to attempt separation of the SDeV particles from the superinfecting reptarenavirus or hartmanivirus particles. Unfortunately, even with varying centrifugation times (18 h and 4 h) we were unable to separate SDAg and reptarenaviruses or hartmaniviruses into different fractions (Figure 6C).

### Transfection of I/1Ki-**Δ** cells with viral glycoproteins induces production of infectious particles

Because superinfection of I/1Ki-Δ cells with both reptarenaviruses and hartmaniviruses resulted in production of infectious SDeV particles, we wanted to study which of the structural proteins are required for particle production. While the envelope of both classical arenaviruses (genus *Mammarenavirus*) and reptarenaviruses comprises both matrix protein (ZP) and spike complexes, the envelope of hartmaniviruses lacks the ZP^29^. Glycoproteins GP1 and GP2, encoded as a glycoprotein precursor (GPC), form the major portion of the spike complex, which in the case of mammarenaviruses and, presumably, hartmaniviruses comprises also a stable signal peptide^29^. We started by transfecting I/1Ki-Δ cells with the GPCs of HISV-1, Puumala virus (PUUV, an orthohantavirus), and UGV-1 (with and without co-transfected ZP). Additionally, we transfected the cells with HBV S-antigen (S-Ag) bearing plasmid. We included PUUV glycoproteins to the experiment, since orthohantaviruses, like mammarenaviruses, are known to induce persistent infection in their rodent hosts and could thus represent a potential helper virus. Additionally, the GPC of orthohantaviruses is similar to that of arenaviruses in the sense that it gives rise to two glycoproteins, Gn and Gc, which form the spike complex^35^. We could demonstrate the expression of glycoproteins using IF staining for all except HBV S-Ag (Figure 7A). We found SDAg to be mostly cytoplasmic, however, many of the non-transfected cells displayed a punctate SDAg reaction in the nucleus. We could not conclude if the expression of viral glycoproteins affects the localization of SDAg.

**Figure 7.**
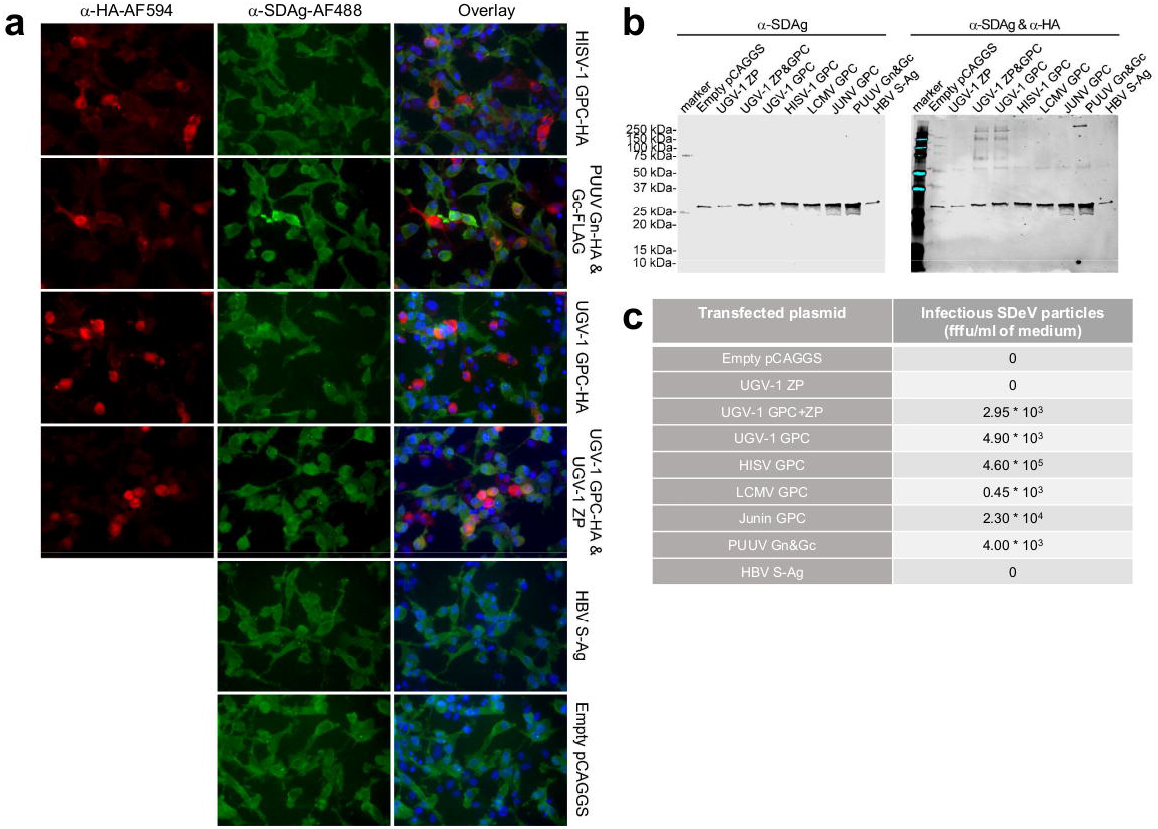
Infectious SDeV particles are formed when I/1Ki-Δ cells are transfected with viral glycoproteins. **a)** I/1Ki-Δ cells transfected with HISV GPC (top row), Puumala virus glycoproteins (PUUV Gn&Gc, second row), UGV-1 GPC (third row), UGV-1 ZP and GPC (fourth row), HBV S-Ag (fifth row), and empty pCAGGS-MCS plasmid (bottom row) were stained for HA-tag (anti-HA 1:4,000 and Alexa Fluor 594-labeled donkey anti-mouse immunoglobulin 1:1,000, left panels, red), SDAg (α-SDAg-AF488, middle panels, green), and Hoechst 33342 was used to visualize the nuclei, the panels on right show an overlay. All images were taken at 400x magnification using Zeiss Axioplan 2 microscope. **b)** Supernatants collected from I/1Ki-Δ cells transfected with empty pCAGGS-MCS plasmid, UGV-1 ZP, UGV-1 GPC and ZP, UGV-1 GPC, HISV-1 GPC, LCMV GPC, JUNV GPC, PUUV glycoproteins, and HBV S-Ag were pelleted by ultracentrifugation and analyzed by western blotting. The left panel shows anti-SDAg staining and the panel on the right shows anti-SDAg and anti-HA staining. **c)** Supernatants collected from I/1Ki-Δ cells transfected with empty pCAGGS-MCS plasmid, UGV-1 ZP, UGV-1 GPC and ZP, UGV-1 GPC, HISV-1 GPC, LCMV GPC, JUNV GPC, PUUV glycoproteins, and HBV S-Ag were titrated on clean I/1Ki cells. The column on the left shows the plasmid used for transfection and the corresponding SDeV titer is shown on the right column.

We then transfected I/1Ki-Δ cells with empty pCAGGS plasmid, UGV-1 ZP, UGV-1 GPC and ZP, UGV-1 GPC, HISV-1 GPC, lymphocytic choriomeningitis virus (LCMV) GPC, Junin virus (JUNV) GPC, PUUV glycoproteins, and HBV S-Ag, and collected the supernatant up to 7 days post transfection. We then used ultracentrifugation to pellet the material secreted from the transfected cells and analyzed the pellets by WB. Transfection of the cells with glycoproteins appeared to enhance the secretion of SDAg (Figure 7B). Some (UGV-1, HISV-1, and PUUV glycoproteins) of the expressed viral glycoproteins were present in high enough amounts in the pellets and could be detected by WB. We wanted to study whether expression of glycoproteins had induced formation of infectious SDeV particles, and infected naïve I/1Ki cells with the collected supernatants. The results show that cells transfected with an empty plasmid, UGV-1 ZP or HBV S-Ag do not produce infectious particles (Figure 7C). However, infectious particles were formed by cells co-transfected with UGV-1 ZP and GPC. Curiously, expression of UGV-1 GPC alone produced higher amounts of infectious particles, indicating that the expression of ZP is not needed for the production of infectious SDeV particles (Figure 7C). Similarly to the superinfection experiments, in which HISV-1 was found as the most effective helper virus, the expression of HISV-1 GPC induced the highest concentration of infectious SDeV particles. Interestingly, the expression of the mammarenavirus and even orthohantavirus GPCs induced production of infectious particles.

## DISCUSSION

Until 2018 the genus *Deltavirus* was represented by a single species, HDV, which was intimately linked with HBV infection. HDV is a satellite virus of HBV which mostly targets the human liver^24^, and hence HDV infection is mostly associated with liver disease. HDV infection affects around 20 million people worldwide^36^, however, it is currently somewhat of a neglected disease due to the fact that HBV vaccination is thought to also protect against HDV infection. The recent discovery of deltaviruses in the absence of hepadnaviral co-infection across a wide range of taxa^5, 6, 28^ provided the first indications that deltaviruses could be much more enigmatic than originally thought. We found SDeV in several tissues of the infected snakes, indicating that the virus has a broad tissue tropism and further suggesting that SDeV does not rely on a hepadnavirus as its helper^6^. Indeed, earlier this year Perez-Vargas and colleagues demonstrated that HDV can form infectious particles utilizing envelope glycoproteins of viruses from various species^27^. We originally found SDeV in co-infection with reptarena- and hartmaniviruses^6^, and wanted to study if these viruses could provide glycoprotein-decorated envelope for the production of infectious SDeV particles.

Here, we generated plasmid constructs bearing the SDeV genome in duplicate and in either genomic or antigenomic orientation, and could demonstrate that transfecting these plasmids into cultured snake, monkey or human cells initiates SDeV replication. These findings imply that SDeV replication itself is not limited to any particular cell type, but would rather be restricted by the envelope borrowed from the co-/superinfecting helper virus. Similarly, HDV infection is likely not restricted to the liver since Perez-Vargas and colleagues demonstrated infectious HDV particle formation in co-infection with various enveloped viruses^27^. By passaging of the cells transfected with SDeV constructs we could demonstrate that, at least in cell culture, deltaviruses can rather easily establish a persistent infection. The persistently infected cell lines allowed us to imitate SDeV infection *in vitro* and to overcome the problems faced in human hepatitis virus research i.e. the lack of a solid cell culture system that allows viral infection and propagation^37^. Moreover, our results not only show that SDeV can establish and maintain a persistent infection *in vitro*, but also indicate that helper virus is not required for persistent infection. HBV-independent persistence of HDV and subsequent rescue by HBV superinfection has been shown in woodchucks^38^, chimpanzees^39^ and mice^40, 41^. HDV can persist in human hepatocytes (in the liver of humanized mice) without its helper virus and potentially be rescued by a later HBV infection^41^. The persistence of SDeV in snake cell lines shown in this study and the persistence of HDV in human hepatocytes^42^ resemble each other in the sense that the accumulation of positive cells appears to rely on cell division rather than a helper virus.

The persistently SDeV infected cell lines enabled us to study the release of SDeV RNA and SDAg in cells superinfected with reptarenaviruses and hartmaniviruses. To our surprise we found I/1Ki-Δ to secrete both SDeV RNA and SDAg even without superinfection. However, clean cells could not be infected using the material released from non-superinfected cells. We also observed that the cellular distribution of SDAg differed between cell lines, ranging from mostly nuclear in liver and heart (V/1Liv and V/2Hz) to mainly cytoplasmic in lung and kidney (V/5Lu and V/1Ki) cell lines. The cellular distribution of HDAg changes during the viral life cycle, the viral ribonucleoproteins are transported back and forth between the nucleus and the cytoplasm, however this transport is influenced by different factors^30^. It could be that some cell lines express proteins capable of triggering re-distribution of SDAg. Such proteins could include e.g. hepadnaviral and lentiviral glycoproteins integrated into the hosts’ genome. In fact, infectious HDV particles can be formed via low level expression of genome-integrated HBV S-Ag^43^. It is tempting to speculate that endogenous viral elements found in animals, plants, and fungi would also contribute to the formation of infectious deltavirus particles and thereby also to their spread. Alternatively, it could be speculated that the observed cellular distribution of SDAg in kidney and lung cells would be due to secretion within vesicles. Supporting the latter hypothesis, we observed that the SDeV RNA and SDAg secreted from I/1Ki-Δ migrates to the same fractions as reptarenaviruses and hartmaniviruses in density gradient ultracentrifugation. By superinfecting I/1Ki-Δ with reptarenaviruses and hartmaniviruses we could induce production of infectious SDeV particles. We could further demonstrate that the expression of glycoproteins alone induces formation of infectious SDeV particles, even though the ZP of arenaviruses is known to contribute to budding of virions^44^. The co-transfection of ZPs did not improve the efficiency of SDeV particle production. Interestingly, hartmanivirus infection or the expression of hartmanivirus glycoprotein seemed to induce the most efficient production of infectious SDeV particles. The dissimilarity could be due to the suggested differences between reptarenavirus (no cytoplasmic tail) and hartmanivirus (cytoplasmic tail with putative late domains) glycoproteins^29^. We could further show that the expression of mammarenavirus (LCMV or JUNV) and hantavirus (PUUV) glycoproteins also induces formation of infectious SDeV particles. We have also applied the same approach for HDV and observed that the expression of arenavirus and hantavirus glycoproteins is sufficient for infectious particle production. Since parallel findings using hepacivirus, flavivirus and vesiculovirus helpers have been recently published by Perez-Vargas and co-authors^27^, we have decided not to include our results regarding HDV in this manuscript. Taken altogether, these findings imply that various deltaviruses would rely on several different helper viruses to complete their life cycle.

These newly described characteristics of deltaviruses raise numerous questions regarding the range of possible helper viruses, and factors contributing to deltavirus-glycoprotein interactions and subsequent infectious particle formation. Further studies will need to address which viruses can act as deltavirus helpers. It is tempting to speculate that deltaviruses would be opportunistic microbes, the exit (or infectious particle formation) of which would rely on persistent, latent, or recurring infections by enveloped viruses. Such infections could include arenaviruses and hantaviruses, both of which cause persistent infection in rodents^45, 46^. In humans, examples of persistent viral infections could include HBV, hepatitis C virus, and human immunodeficiency virus, the first two of which have already been demonstrated to induce the formation of infectious HDV particles^27^. Additionally, recurring infections such as those caused by ortho-, paramyxo- or coronaviruses, or latent infections caused by e.g. herpesviruses could contribute to the spread of deltaviruses. It seems thus fair to speculate that HDV could present merely the tip of an iceberg in terms of human deltavirus infections. This theory would be supported by the detection of HDAg in the absence of HBV from the salivary glands of Sjögren’s syndrome patients^47^. Although it is unclear if and how the HDAg expression (or viral replication) directly contributes to the development of Sjögren’s syndrome, it appears that deltaviruses could be the underlying cause or agents able to exacerbate disease in both animals and humans. The common ancestor of deltaviruses could be found among viroids of higher plants, since they show the highest similarities with HDV^4^. Since a deltavirus was already identified in termites^28^, one could hypothesize that the deltavirus ancestor was transmitted to Animalia from plants. Indeed, Bogaert and colleagues showed the presence of viroids in aphids feeding on infected plants^48^ which could support the above hypothesis.

Deltaviruses are likely widespread both worldwide and across different taxa. So far novel deltaviruses have been found in snakes, birds, fish, amphibians, and invertebrates^5, 6, 28^. These findings add to the previously limited knowledge about the origin and evolution of HDV^7^. Also the findings on the infidelity of HDV to HBV^27^ indicate that many new doors have lately been opened in the field of deltavirus research.

## MATERIALS AND METHODS

### Cell lines and viruses

We used the following established cell lines: human hepatocellular carcinoma, Hep G2 (American type culture collection, ATCC); African green monkey kidney, Vero E6 (ATCC); human embryonic kidney, HEK293FT (Thermo Fisher Scientific); human lung carcinoma, A549 (ATCC). *Boa constrictor* kidney cell line, I/1Ki described in Hetzel et al., 2013^32^. Additionally, we established the following cell lines by applying techniques described in Hetzel et al., 2013^32^; *B. constrictor* kidney, V/1Ki; *B. constrictor* liver, V/1Liv; *B. constrictor* lung, V/5Lu; and *B. constrictor* heart, V/2Hz.

For maintaining the cultured mammalian and I/1Ki cell lines we used Minimal Essential Medium Eagle (MEM) supplemented with 10% fetal bovine serum, 200 mM L-glutamine, 100 μg/ml of streptomycin, and 100 U/ml of penicillin, while for the other snake cell lines we used Dulbecco’s Modified Eagle Media (DMEM) – high glucose supplemented with 15% fetal bovine serum, 200 nM L-Alanyl L-glutamine, 100 μg/ml of streptomycin, and 100 U/ml of penicillin. We kept the cells in incubators with 5% CO_2_ and at 30 °C or 37 °C.

To obtain cell lines persistently infected with SDeV, we passaged the cells (culturing conditions as described above) transfected with plasmid bearing two copies of the SDeV genome (described below) until we obtained a confluent 175-cm^2^ flask and were able to prepare ampoules for storage. The following permanently infected cell lines were generated: I/1Ki-Δ, V/1Ki-Δ, V/1Liv-Δ, V/2Hz-Δ, and V/5Lu-Δ.

For superinfection studies we used two reptarenaviruses: University of Helsinki virus-2 (UHV-2^29^) and University of Giessen virus-1 (UGV-1^33^), and one hartmanivirus, Haartman Institute Snake virus-1 (HISV-1^29^).

### Cloning, plasmids and recombinant protein expression

We ordered a synthetic gene from Gene Universal bearing the snake deltavirus genome (MH988742.1) in duplicate (starting from residue 216 and ending at 215 i.e. exactly two copies of the genome) with T7 promoter (5’-TAATACGACTCACTATAGG-3’) after the SDeV genome, and EcoRV restriction sites at both ends. We followed the manufacturer’s protocols throughout the cloning. We used FastDigest EcoRV (ThermoFisher Scientific), agarose gel electrophoresis and the GeneJET Gel extraction kit (ThermoFisher Scientific) to purify the synthetic insert. For subcloning the synthetic insert to pCAGGS/MCS as described in Niwa et al., 1991^49^, we used FastDigest EcoRI and XhoI (both ThermoFisher Scientific) to linearize the plasmid, T4 DNA ligase (ThermoFisher Scientific) to blunt the 5’ and 3’ overhangs, and the GeneJET Gel extraction kit (ThermoFisher Scientific) to purify the plasmid after agarose gel electrophoresis. We ligated the insert to pCAGGS/MCS using T4 DNA ligase (ThermoFisher Scientific), transformed chemically competent *E. coli* (DH5α strain) with standard methods, plated the bacteria on Luria-Broth agar plates with 100 μg/ml of ampicillin, picked single colonies after overnight (O/N) cultivation at 37 °C into 5 ml of 2xYT medium (16 g/l tryptone, 10 g/l yeast extract, 5 g/l NaCl), prepared minipreps from 2 ml of O/N cultivation at 37 °C using the GeneJET Plasmid Miniprep Kit (ThermoFisher Scientific), checked for the presence of insert using restriction digest and agarose gel electrophoresis, Sanger sequenced (DNA Sequencing and Genomic Laboratory, Institute of Biotechnology, University of Helsinki) the plasmids to obtain clones with insert in genomic and antigenomic orientation, and used ZymoPURE II Plasmid Maxiprep Kit (Zymo Research) to obtain plasmid stocks for transfection.

For recombinant expression of HBV S-Ag, we ordered a synthetic gene (from GeneUniversal) covering the residues 2848-3215 and 1-835 of HBV (JX079936.1), which encode large, middle, and small S-Ag, with 5’ EcoRI and 3’ XhoI restriction sites, subcloned the insert into pCAGGS/MCS^49^ and prepared plasmid stocks as described above. We also ordered codon-optimized (for human) synthetic genes (from GeneUniversal) based on UGV-1 ZP (AKN10693.1), with 5’ EcoRI and 3’ XhoI restriction sites, and cloned the gene to pCAGGS/MCS for expression as described above. For cloning of UGV-1 GPC and HISV GPC we used RT-PCR amplification (reverse transcription with RevertAid Reverse Transcriptase, ThermoFisher Scientific; PCR with Phusion Flash High-Fidelity PCR Master Mix, ThermoFisher Scientific) from RNA extracted (by GeneJET RNA Extraction Kit, ThermoFisher Scientific) with the following primers: UGV-GPC-fwd 5’-AAAGAATTCATGGCAGGTCACCTCAACCG-3’, UGV-GPC-rev 5’-TTTATGCATCCCCGTCTCACCCAGTTGC-3’, HISV-GPC-fwd 5’-AAAGAATTCATGGGGGCACTTGTGTCC-3’ and HISV-GPC-rev 5’-GGAGGTACCCCGTATTTTTCAATGGGACA-3’ to generate inserts, purified the inserts using the GeneJET PCR Purification Kit (ThermoFisher Scientific), restriction digested the inserts with FastDigest EcoRI and SmaI (ThermoFisher Scientific) for UGV-1 GPC and FastDigest EcoRI and KpnI (ThermoFisher Scientific) for HISV-1 GPC, purified the inserts as described above, ligated with Thermo Selective Alkaline Phosphatase treated (ThermoFisher Scientific) pCAGGS-HA^50^, linearized with the respective restriction enzymes, and prepared the plasmid stocks as described above. Lymphocytic choriomeningitis virus (LCMV) and Junin virus (JUNV) GPCs were amplified using primers 5’-AATTCAATTGACCATGGGTCAGATTGTG-3’ and 5’-AATTCCCGGGGCGTCTTTTCCAGAC-3 (LCMV GPC) or 5’-AATTGAGCTCACCATGGGGCAGTTCATT-3’ and 5’-AATTCCCGGGGTGTCCTCTACGCCA-3’ (JUNV GPC) and cloned using EcoRI/MfeI and SmaI (LCMV GPC); or SacI and SmaI (JUNV GPC) into pCAGGS-HA. Plasmids were verified by sequencing. For cloning of Puumala orthohantavirus (PUUV) Gn (residues 1-658 of CCH22848.1) and Gc (residues 637-1148 of CCH22848.1) we used the following primers: PUUV-Gn-fwd 5’-AATAGAATTCATGGGAAAGTCCAGCCCCGTGT-3’, PUUV-Gn-rev 5’-TCCCGGGTGCGCTGGCGGCCCACA-3’, PUUV-Gc-fwd 5’-AAGAGAATTCATGTTCTTCGTGGGCCT-3’, PUUV-Gc-rev 5’-ATTCCCGGGCTTGTGCTCCTTC-3’ to PCR amplify (Phusion Flash High-Fidelity PCR Master Mix, ThermoFisher Scientific) the inserts from codon optimized PUUV GPC described in Iheozor-Ejiofor et al., 2016^51^, ligated the inserts employing EcoRI and SmaI restriction sites to both pCAGGS-HA and pCAGGS-FLAG^52^ and prepared plasmid stocks as described above.

### Affinity purification and labelling of antibodies

To generate affinity purified anti-UHV NP and anti-SDAg antibodies, we coupled baculovirus expressed recombinant UHV NP^53^ and *E. coli* expressed SDAg^6^ to CNBr-Activated Sepharose 4 Fast Flow (GE Healthcare Life Sciences) following the manufacturer’s protocol. We used a protocol described in Korzyukov et al., 2016^54^ to affinity purify the antibodies. After dialysis and concentration of the antibodies we labelled the affinity purified fractions using Alexa Fluor 488 TFP ester (ThermoFisher Scientific) or Alexa Fluor 594 NHS Ester (ThermoFisher Scientific) following the manufacturer’s recommendation. The labelled antibodies were purified by passing through Disposable PD 10 Desalting Columns (GE Healthcare Life Sciences), concentrated using Amicon Ultra-15 Centrifugal Filter Units (Millipore) with 50K nominal molecular weight cutoff, mixed with Glycerol (final concentration 50% v/v), and kept at −20 °C for short term and at −80 °C for long term storage. To determine optimal dilutions, we titrated the antibodies on clean and infected I/1Ki-Δ cells (SDeV with and without reptarenavirus infection), the antibodies generated are (dilution range in brackets): anti-UHV NP-AF488 (1:200-1:400), anti-UHV NP-AF594 (1:200-1:400), anti-SDAg-AF488 (1:400-1:800), and anti-SDAg-AF594 (1:400-1:800).

### SDS-PAGE and immunoblotting

We performed SDS-PAGE (self-prepared gels and 4–20% Mini-PROTEAN® TGX gels from Bio-Rad) and immunoblotting using methods described in Korzyukov et al., 2016^54^. The antibody dilutions used were 1:10,000 to 1:20,000 for the rabbit anti-SDAg pAb^6^, 1:4,000 for the mouse anti HA-tag mAb (AE008, ABclonal), 1:10,000 for AlexaFluor 680 donkey anti-rabbit (IgG) and anti-mouse (IgG) (ThermoFisher Scientific), and 1:10,000 for IRDye 800CW donkey anti-rabbit (IgG) and anti-mouse (IgG) (LI-COR Biosciences). We used the Odyssey Infrared Imaging System (LI-COR Biosciences) for recording the results.

### Transfection of cultured cells

For transfection of *B. constrictor* cell lines we used Lipofectamine 2000 and for transfection of mammalian cells we used Fugene HD (Promega). When using Lipofectamine 2000 we prepared the transfection mixes diluting the plasmid stock in OptiMEM (ThermoFisher Scientific) to yield 500 ng/50 μl, mixed 2.75-3.0 μl of Lipofectamine 2000 in 47 μl of OptiMEM (ThermoFisher Scientific), combined the two mixtures by pipetting up and down, and allowed the complexes to form 15-30 min at room temperature (RT). When using Fugene HD we prepared the plasmid solution as above, added 1.75 μl of Fugene HD, mixed by pipetting, and allowed the complexes to form (15-30 min at RT). The above recipes were used for transfecting 2 cm^2^ of cells, and scaled up based on the desired cell surface area. After preparing the transfection reagent-plasmid complexes, we detached 80-90% confluent cell layers using Gibco Trypsin-EDTA (0.25%, ThermoFisher Scientific), pelleted the cells by centrifugation (3-4 min at 500g), re-suspended the cells into fully conditioned cell culture medium (described above) to yield a cell density of approximately 2 cm^2^/ml (e.g. cells from a confluent 75 cm^2^ bottle would be re-suspended in 37.5 ml), mixed 1 ml of cell suspension with the transfection mix by pipetting, incubated for 15-30 min at RT, plated the cells, and replaced the medium after 6 h incubation.

### Immunofluorescence (IF) staining

We used black 96-well plates (PerkinElmer) or 13 mm coverslips to grow the cells for IF staining. For collagen coating, we incubated the coverslips or plates O/N at +4 °C with 0.1 mg/ml of collagen I from rat tail (BD Biosciences) in 0.25% acetic acid. For fixing the cells we removed the culture medium, added 4% paraformaldehyde (PFA) in PBS (pH 7.4), incubated for 10 min at RT, washed once with TBS (50 mM Tris, 150 mM NaCl, pH 7.4), permeabilized and blocked (0.25% Triton X-100 [Sigma Aldrich], 3% bovine serum albumin [BSA, ThermoFisher Scientific] in TBS) for 5-10 min at RT and washed once with TBS. For IF staining we incubated the cells with the primary antibodies diluted in TBS with 0.5% BSA for 60-90 min at RT, washed three times with TBS, incubated 45 min with the secondary antibody diluted in TBS with 0.5% BSA, washed three times with TBS, once with Hoechst 33342 diluted in TBS, once with TBS, twice with MilliQ water (Millipore), and mounted the coverslips with Prolong Gold Antifade Mountant (ThermoFisher Scientific) or added 50 μl/well of 90% glycerol, 25 mM Tris-HCl, pH 8.5, for the 96-well plates. For primary antibodies we used the following dilutions: 1:7,500 for anti-SDAg^6^, 1:2,500 for anti-UHV NP-C^53^, 1:2,500 for anti-HISV NP-C^29^, 1:250 for monoclonal anti-HA (ABclonal); and for secondary antibodies: 1:1,000 for Alexa Fluor 488- or 594-labeled donkey anti-rabbit immunoglobulin (ThermoFisher Scientific) and 1:1,000 for AlexaFluor 488- or 594-labeled donkey anti-mouse immunoglobulin (ThermoFisher Scientific).

### Virus purification

We used ultracentrifugation to pellet viruses produced by reptarena- and hartmanivirus superinfected I/1Ki-Δ cells (*B. constrictor* kidney I/1Ki permanently infected with snake deltavirus). Briefly, we collected the supernatants up to 14 days post infection (dpi), cleared by centrifugation (30 min, 4200g, +4 °C), filtered through a 0.45 μm syringe filter (Millipore), placed into either 25×89 mm (for SW28 rotor) or 14×89 mm (for SW41 rotor) Ultra-Clear tubes (Beckman Coulter), underlaid with 3 ml (for SW28 rotor) or 1 ml (for SW41 rotor) of 25% (w/v) sucrose (in TBS) using a thin needle, performed ultracentrifugation (27,000 rpm, 1.5-2 h, +4 °C, for both SW28 and SW41 rotor), poured off the supernatant and sucrose cushion, and resuspended the pellet in TBS. We used the same protocol for concentrating the supernatants collected from transfected I/1Ki-Δ cells. For density gradient ultracentrifugation we used Gradient Master (BioComp) to prepare 10-70% linear sucrose gradients (in TBS) in 14×89 mm (for SW41 rotor) Ultra-Clear tubes (Beckman Coulter), following the manufacturer’s protocol. We loaded the viruses concentrated as described above on top of the gradient, performed ultracentrifugation (40,000 rpm, either 4 h or 18 h, +4 °C, SW41 rotor, Beckman Coulter), and collected approximately 600 μl fractions by puncturing the tubes with a thin (23G or 25G) needle.

### Virus titration

To determine if superinfections or transfections with viral glycoproteins had induced formation of infectious SDeV particles, we performed 10-fold dilution series of the supernatants, and used the diluted supernatants to inoculate uninfected I/1Ki or Vero E6 cells. At four to six dpi the cells were fixed and stained for SDAg as described above. The number of fluorescent focus-forming units (fffu) were determined by enumerating the number of SDAg positive cells using fluorescence microscopy. Each dilution was represented by two or three parallel wells, and, when possible, two consecutive dilutions were used for calculating the number of fffus in the original sample.

## DATA AVAILABILITY

All data generated or analysed in this study are included in this published article (and its supplementary information files).

## ACKNOWLEDGEMENTS

We are grateful for Prof. Juan Carlos de la Torre for kindly providing the pCAGGS-HA and pCAGGS-FLAG plasmids used in this study. This study was supported by the following grants: Academy of Finland (JH, grant numbers: 1308613 and 1314119).

## AUTHOR CONTRIBUTIONS

Conception/design of the work: Jussi Hepojoki

Data collection: Leonora Szirovicza, Jussi Hepojoki

Data analysis and interpretation: Leonora Szirovicza, Jussi Hepojoki

Resources: Jussi Hepojoki, Luis Martinez-Sobrido

Funding acquisition: Jussi Hepojoki

Drafting the article: Leonora Szirovicza, Jussi Hepojoki

Critical revision of the article: Leonora Szirovicza, Udo Hetzel, Anja Kipar, Olli Vapalahti, Luis Martinez-Sobrido, Jussi Hepojoki

Final approval of the version to be published: Leonora Szirovicza, Udo Hetzel, Anja Kipar, Luis Martinez-Sobrido, Olli Vapalahti, Jussi Hepojoki

## ADDITIONAL INFORMATION

### Materials & Correspondence

Correspondence and requests for materials should be addressed to Leonora Szirovicza;

## REFERENCES

1. Flores, R., Owens, R. A. & Taylor, J. Pathogenesis by subviral agents: viroids and hepatitis delta virus. Curr. Opin. Virol. 17, 87–94 (2016).

2. Diener, T. O. Potato spindle tuber “virus”. IV. A replicating, low molecular weight RNA. Virology 45, 411–438 (1971).

3. Rizzetto, M. et al. Immunofluorescence detection of new antigen-antibody system (delta/anti-delta) associated to hepatitis B virus in liver and in serum of HBsAg carriers. Gut 18, 997–1003 (1977).

4. Flores, R., Ruiz-Ruiz, S. & Serra, P. Viroids and hepatitis delta virus. Semin. Liver Dis. 32, 201–210 (2012).

5. Wille, M. et al. A Divergent Hepatitis D-Like Agent in Birds. Viruses 10, 720 (2018).

6. Hetzel, U. et al. Identification of a Novel Deltavirus in Boa Constrictors. MBio 10, e00014–00019 (2019).

7. Littlejohn, M., Locarnini, S. & Yuen, L. Origins and Evolution of Hepatitis B Virus and Hepatitis D Virus. Cold Spring Harb. Perspect. Med. 6, a021360 (2016).

8. Magnius, L. et al. ICTV Virus Taxonomy Profile: Deltavirus. J. Gen. Virol. 99, 1565–1566 (2018).

9. Wang, K. S. et al. Structure, sequence and expression of the hepatitis delta (delta) viral genome. Nature 323, 508–514 (1986).

10. Kos, A., Dijkema, R., Arnberg, A. C., van der Meide, P. H. & Schellekens, H. The hepatitis delta (δ) virus possesses a circular RNA. Nature 323, 558–560 (1986).

11. Chen, P. et al. Structure and replication of the genome of the hepatitis δ virus. Proc. Natl. Acad. Sci. USA 83, 8774–8778 (1986).

12. Kuo, M. Y., Sharmeen, L., Dinter-Gottlieb, G. & J., T. Characterization of self-cleaving RNA sequences on the genome and antigenome of human hepatitis delta virus. J. Virol. 62, 4439–4444 (1988).

13. Sharmeen, L., Kuo, M. Y. & Taylor, J. Self-ligating RNA sequences on the antigenome of human hepatitis delta virus. J. Virol. 63, 1428–1430 (1989).

14. Reid, C. E. & Lazinski, D. W. A host-specific function is required for ligation of a wide variety of ribozyme-processed RNAs. Proc. Natl. Acad. Sci. USA 97, 424–429 (2000).

15. Sureau, C., Guerra, B. & Lanford, R. E. Role of the large hepatitis B virus envelope protein in infectivity of the hepatitis delta virion. J. Virol. 67, 366–372 (1993).

16. Rizzetto, M. et al. Delta agent: association of delta antigen with hepatitis B surface antigen and RNA in serum of delta-infected chimpanzees. Proc. Natl. Acad. Sci. USA 77, 6124–6128 (1980).

17. Kuo, M. Y., Chao, M. & J., T. Initiation of replication of the human hepatitis delta virus genome from cloned DNA: role of delta antigen. J. Virol. 63, 1945–1950 (1989).

18. Chang, J., Nie, X., Chang, H. E., Han, Z. & Taylor, J. Transcription of hepatitis delta virus RNA by RNA polymerase II. J. Virol. 82, 1118–1127 (2008).

19. Chao, M., Hsieh, S. & Taylor, J. Role of two forms of hepatitis delta virus antigen: evidence for a mechanism of self-limiting genome replication. J. Virol. 64, 5066–5069 (1990).

20. Luo, G. X. et al. A specific base transition occurs on replicating hepatitis delta virus RNA. J. Virol. 64, 1021–1027 (1990).

21. Chang, F., Chen, P., Tu, S., Wang, C. & Chen, D. The large form of hepatitis δ antigen is crucial for assembly of hepatitis δ virus. Proc. Natl. Acad. 88, 8490–8494 (1991).

22. Krogsgaard, K. et al. Delta-infection and suppression of hepatitis B virus replication in chronic HBsAg carriers. Hepatology 7, 42–45 (1987).

23. Huang, C. R. & Lo, S. J. Hepatitis D virus infection, replication and cross-talk with the hepatitis B virus. World J. Gastroenterol. 20, 14589–14597 (2014).

24. Farci, P. & Niro, G. A. Clinical features of hepatitis D. Semin. Liver Dis. 32, 228–236 (2012).

25. Negro, F. Hepatitis D virus coinfection and superinfection. Cold Spring Harb. Perspect. Med. 4, a021550 (2014).

26. Le Gal, F. et al. Eighth major clade for hepatitis delta virus. Emerg. Infect. Dis. 12, 1447–1450 (2006).

27. Perez-Vargas, J. et al. Enveloped viruses distinct from HBV induce dissemination of hepatitis D virus in vivo. Nat. Commun. 10, 2098 (2019).

28. Chang, W.-S., et al. Novel hepatitis D-like agents in vertebrates and invertebrates. Preprint at: https://www.biorxiv.org/content/10.1101/539924v1.

29. Hepojoki, J. et al. Characterization of Haartman Institute snake virus-1 (HISV-1) and HISV-like viruses-The representatives of genus Hartmanivirus, family Arenaviridae. PLoS Pathog 14, e1007415 (2018).

30. Tavanez, J. P. et al. Hepatitis delta virus ribonucleoproteins shuttle between the nucleus and the cytoplasm. RNA 8, 637–646 (2002).

31. Canese, M. G. et al. An ultrastructural and immunohistochemical study on the delta antigen associated with the hepatitis B virus. J. Pathol. 128, 169–175 (1979).

32. Hetzel, U. et al. Isolation, identification, and characterization of novel arenaviruses, the etiological agents of boid inclusion body disease. J. Virol. 87, 10918–10935 (2013).

33. Hepojoki, J. et al. Arenavirus coinfections are common in snakes with boid inclusion body disease. J. Virol. 89, 8657–8660 (2015).

34. Dervas, E. et al. Nidovirus-associated proliferative pneumonia in the green tree python (Morelia viridis). J. Virol. 91, e00718–00717 (2017).

35. Hepojoki, J., Strandin, T., Vaheri, A. & Lankinen, H. Interactions and oligomerization of hantavirus glycoproteins. J. Virol. 84, 227–242 (2010).

36. Cunha, C., Tavanez, J. P. & Gudima, S. Hepatitis delta virus: A fascinating and neglected pathogen. World J. Virol. 4, 313–322 (2015).

37. Sureau, C. The use of hepatocytes to investigate HDV infection: the HDV/HepaRG model. Methods Mol. Biol. 640, 463–473 (2010).

38. Netter, H. J., Gerin, J. L., Tennant, B. C. & Taylor, J. M. Apparent helper-independent infection of woodchucks by hepatitis delta virus and subsequent rescue with woodchuck hepatitis virus. J. Virol. 68, 5344–5350 (1994).

39. Smedile, A. et al. Hepatitis D viremia following orthotopic liver transplantation involves a typical HDV virion with a hepatitis B surface antigen envelope. Hepatology 27, 1723–1729 (1998).

40. Polo, J. M. et al. Transgenic mice support replication of hepatitis delta virus RNA in multiple tissues, particularly in skeletal muscle. J. Virol. 69, 4880–4887 (1995).

41. Giersch, K. et al. Persistent hepatitis D virus mono-infection in humanized mice is efficiently converted by hepatitis B virus to a productive co-infection. J. Hepatol. 60, 538–544 (2014).

42. Giersch, K. et al. Hepatitis delta virus persists during liver regeneration and is amplified through cell division both in vitro and in vivo. Gut 68, 150–157 (2019).

43. Freitas, N., Cunha, C., Menne, S. & Gudima, S. O. Envelope proteins derived from naturally integrated hepatitis B virus DNA support assembly and release of infectious hepatitis delta virus particles. J. Virol. 88, 5742–5754 (2014).

44. Urata, S. & Yasuda, J. Molecular mechanism of arenavirus assembly and budding. Viruses 4, 2049–2079 (2012).

45. Rawls, W. E., Chan, M. A. & Gee, S. R. Mechanisms of persistence in arenavirus infections: a brief review. Can. J. Microbiol. 27, 568–574 (1981).

46. Meyer, B. J. & Schmaljohn, C. S. Persistent hantavirus infections: characteristics and mechanisms. Trends Microbiol. 8 (2000).

47. Weller, M. L. et al. Hepatitis delta virus detected in salivary glands of Sjögren’s syndrome patients and recapitulates a Sjögren’s syndrome-like phenotype in vivo. Pathog. Immun. 1, 12–40 (2016).

48. Van Bogaert, N., De Jonghe, K., Van Damme, E. J., Maes, M. & Smagghe, G. Quantitation and localization of pospiviroids in aphids. J. Virol. Methods 211, 51–54 (2015).

49. Niwa, H., Yamamura, K. & Miyazaki, J. Efficient selection for high-expression transfectants with a novel eukaryotic vector. Gene 108, 193–199 (1991).

50. Martinez-Sobrido, L., Giannakas, P., Cubitt, B., Garcia-Sastre, A. & de la Torre, J. C. Differential inhibition of type I interferon induction by arenavirus nucleoproteins. J. Virol. 81, 12696–12703 (2007).

51. Paneth Iheozor-Ejiofor, R., et al. Vaccinia virus-free rescue of fluorescent replication-defective vesicular stomatitis virus and pseudotyping with Puumala virus glycoproteins for use in neutralization tests. J. Gen. Virol. 97, 1052–1059 (2016).

52. Ortiz-Riano, E., Cheng, B. Y., de la Torre, J. C. & Martinez-Sobrido, L. The C-terminal region of lymphocytic choriomeningitis virus nucleoprotein contains distinct and segregable functional domains involved in NP-Z interaction and counteraction of the type I interferon response. J. Virol. 85, 13038–13048 (2011).

53. Hepojoki, J. et al. Replication of boid inclusion body disease-associated arenaviruses is temperature sensitive in both boid and mammalian cells. J. Virol. 89, 1119–1128 (2015).

54. Korzyukov, Y., Hetzel, U., Kipar, A., Vapalahti, O. & Hepojoki, J. Generation of Anti-Boa Immunoglobulin Antibodies for Serodiagnostic Applications, and Their Use to Detect Anti-Reptarenavirus Antibodies in Boa Constrictor. PLoS One 11, e0158417 (2016).

